# Hyperparameter optimisation in differential evolution using Summed Local Difference Strings, a rugged but easily calculated landscape for combinatorial search problems

**DOI:** 10.1101/2023.07.11.548503

**Authors:** Husanbir Singh Pannu, Douglas B. Kell

## Abstract

We analyse the effectiveness of differential evolution hyperparameters in large-scale search problems, i.e. those with very many variables or vector elements, using a novel objective function that is easily calculated from the vector/string itself. The objective function is simply the sum of the differences between adjacent elements. For both binary and real-valued elements whose smallest and largest values are min and max in a vector of length N, the value of the objective function ranges between 0 and *(N-1) × (max-min)* and can thus easily be normalised if desired. This provides for a conveniently rugged landscape. Using this we assess how effectively search varies with both the values of fixed hyperparameters for Differential Evolution and the string length. String length, population size and generations for computational iterations have been studied. Finally, a neural network is trained by systematically varying three hyper-parameters, viz population (NP), mutation factor (F) and crossover rate (CR), and two output target variables are collected (a) median and (b) maximum cost function values from 10-trial experiments. This neural system is then tested on an extended range of data points generated by varying the three parameters on a finer scale to predict both *median* and *maximum* function costs. The results obtained from the machine learning model have been validated with actual runs using Pearson’s coefficient to justify the reliability to motivate the use of machine learning techniques over grid search for hyper-parameter search for numerical optimisation algorithms. The performance has also been compared with SMAC3 and OPTUNA in addition to grid search and random search.

## 1. Introduction

The tunably rugged fitness landscape reflects the intuition that combinatorial search problems can be seen in terms of a ‘landscape’ containing valleys and hills [1]. The NK model invented by Stuart Kauffman [2] can be adjusted by changing N (string length) and K to define the ruggedness level of the flexible landscape as shown in Figure To explain the search of most rugged string in the NK-landscape, a random three letter string has been considered (JIE) in Fig 1(A). In the consecutive iterations, various single letter alterations have been considered in each iteration and cumulative distance of consecutive letters has been used for the cost function (Fig 1(B)). Furthermore, the connected graph shows potential search paths in pursuit of extreme points (highest peak and deepest valley) within the combinatorial search space of string sequences. Darkness in colours in Fig. 1(C) signifies higher cost functions and finally in Fig. 1(D) this graph is one of the various tracks in complex NK-landscape showing the challenge of the optimization algorithm.

**Fig 1.**
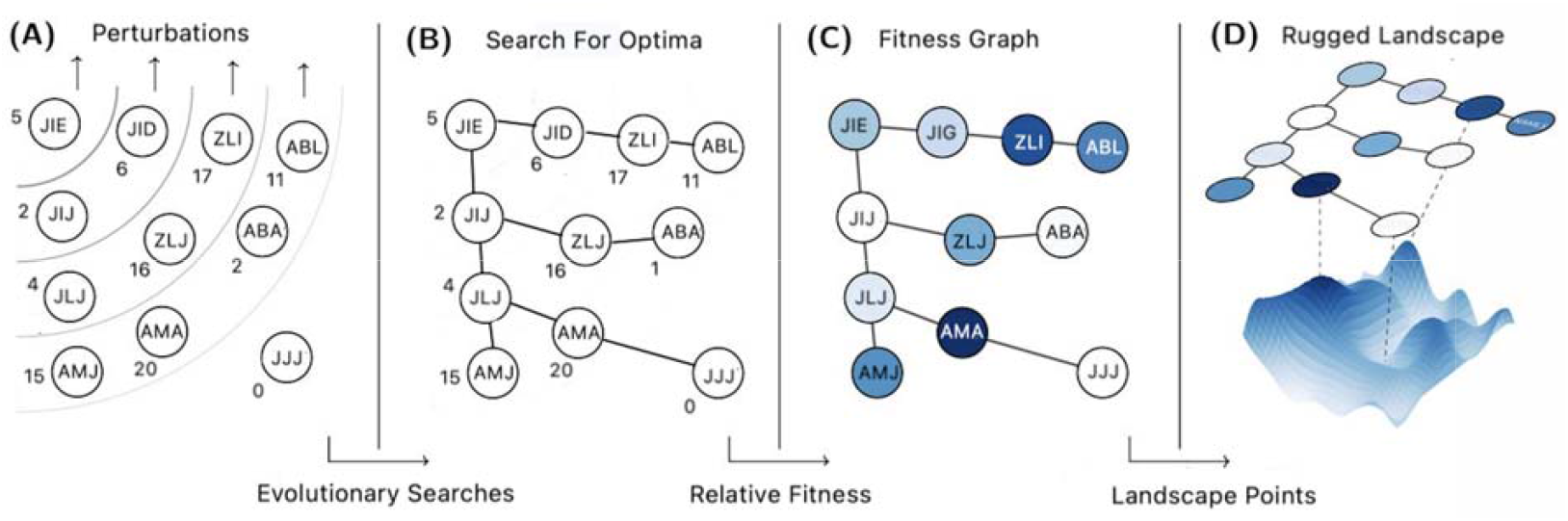
(A) Combination space of various example string sequences of length 3. The digits represent the cumulative inter-letter distances of the string; the sequences along the arrows show the alterations of letters within consecutive strings to signify perturbations (B) Graphical rendering of potential paths to search for the global extrema (peak or valley) in the combinatorial string space (C) Relative colouring for visual comparison, darker means bigger cost function value i.e. higher cost function value (D) The graphs is one of the search possibility in the high NK-landscape showing the complexity of the search of extreme points through an algorithm.

Combinatorial search problems are common in both theoretical and applied sciences such as optimisations, biology and complex evolutionary systems. As an example, in a business organisation, an agent may be searching a landscape of business opportunities. The valleys and hills represent the losses and profits. The journey through the landscape associates the decisions of the organisation whilst altering the structure of the organisation and modifying the products and services. All of the underlying processes interact in a complex evolutionary fashion and affect the cash flow and thus profit [3]. Most scientific problems can in fact be cast as combinatorial search or optimisation problems [4]. The kinds of optimisation in which we are here interested thus consist of a ‘search space’ or landscape in which a variety of inputs can be combined in potentially complex and nonlinear ways to lead to an output or objective function. Because the number of combinations always scales exponentially with the number of variables, the search spaces can easily be made to be far beyond any kind of exhaustive search (other than for small numbers of variables [5]), whether the search is computational or experimental. Heuristic methods, in which we seek to understand, simulate and navigate the landscape intelligently, are therefore appropriate. Among these, evolutionary algorithms of various kinds are pre-eminent [6]. Equally, because of the vastness of the search spaces, a variety of attempts have been made to create them *in silico*, using a more-or-less complex function of the inputs to calculate the output at that position in the search space. The idea of such strategies is that (notwithstanding that there is ‘no free lunch’ [7-11]) an algorithm of interest may be assessed in competition with others [12], or its hyperparameters tuned to effect the most rapid searches. In evolutionary computing, three of these hyperparameters are the population size, the mutation rate, and the crossover rate [13]. In some cases, these hyperparameters can have very considerable effects on the efficiency of a search (e.g. [14]).

Well-known fitness functions used for creating landscapes of this type, often tuned to be ‘deceptive’ to evolutionary algorithms [15,16], include NK [2,17,18] (and variants such as NKp [19]), OneMax/max-ones [20,21], and the royal road [22-24]. Max-ones is especially easy to understand, since each variable is cast as an element of a (binary) string or vector of length N, and the fitness is simply the total number of elements containing a 1, with the maximum possible fitness obviously being N. That said, max-ones yields reasonably easily to evolutionary algorithms [25] as the crossover operator allows such algorithms more-or-less easily to combine building blocks (schemata [13]) successfully, not least because the ‘fitness’ of any element of a string is not context-sensitive.

We here develop and exploit a simple, related objective function in which the objective is not to maximise each element but to maximise *the sum of the differences between adjacent elements*. This is very easily calculated, allowing rapid assessment of different search algorithms. It provides for a landscape that is locally smooth but globally very rugged. Here the contribution to the overall fitness of any element of the string is absolutely context-sensitive. Thus, the problem is to find the most rugged string of length *n* using differential evolution which has three tuning parameters, viz. NP (population size of search particles), F (mutation factor), CR (crossover rate). The objective function for the hyperparameter optimisation for summed local difference strings has been defined with the following examples.

**Example 1:** String = “AAAA” has summed differences |A-A|+|A-A|+|A-A| = 0. String “ABCD” has summed differences |A-B|+|B-C|+|C-D| = 3. String “AZAZ” or “ZAZA” has |A-Z|+|A-Z|+|A-Z| = 75.

**Example 2:** The maximum ruggedness value of a string of length n is *(n-1) × (U-L)* where U and L are the maximum and minimum values of the alphabet values. In example 1 U = Z = 26 and L = A = 1 and n=4, thus (4-1) × (26-1) = 75.

Even for binary strings this makes the problem much harder than max-ones. In addition, two very different (in fact maximally different) strings of length N have the same, maximum fitness, viz 101010…1010101 and 010101…0101010 of (N-1). We later also consider real-valued (integer) strings, such that the problem difficulty can be varied not only by varying the string length but by varying the number of allowable values in each position. Of the many variants of evolutionary algorithm, we focus on differential evolution, as originated by Storn and Price [26,27] and reviewed e.g. in [28-34], as it seems to be highly effective in solving a wide range of problems.

Objective function and machine learning models need to be optimised according to the data distribution in order to find the best representative generalisation. Hyper-parameter tuning determines the best combination of free variables so that validation set yields the best performance. Hyperparameters have been tuned either by manual hit-and-trial, or through grid search, which involves systematically trying all possible combinations with a specified linear spacing. But both of these methods are limited to the human imagination or time duration to attempt grid search on a given level of granularity. Thus, an automatic machine learning based hyperparameter value optimisation has been proposed (e.g. [34-36]).

The paper is organised as follows: the first section is about the introduction of the rugged fitness landscape and local difference strings, Section 2 is a literature review, Section 3 covers the differential evolution and machine learning techniques used, Section 4 is about the results and discussion, Section 5 is the conclusion and future scope.

## 2. Literature Review

### 2.1 Differential Evolution Variants

In [37], a variant of DE using asymptotic termination based on the average differential of the cost function values has been proposed. The second modification is a new search for a critical parameter which helps to explore the search space. In [38], an ensemble of control parameters and mutation methods for DE has been proposed while considering the dynamic mutation strategies and set of values for control parameters. In [39], monkey king differential evolution using a multi-trial vector has been studied. The relation between exploitation and exploration depends upon control parameter and evolution strategy. To enhance the performance, multiple evolution strategies have been considered to generate multi-trial vectors. In [40], an ensemble of DE using multi-population approach and three distinct mutation methods, “rand/1”, “current-to-rand/1”, “current-to-pbest/1”. CEC 2005 benchmark functions have been used for performance evaluation.

### 2.2 Hyper-parameter Tuning

In [41] an advanced hyper-parameter tuning technique ‘Optuna’ has been proposed. It allows API for dynamic user interaction, efficient search and pruning options, and easy to implement features. An automatic parameter tuner for sparse Bayesian learning has been proposed in [42]. The empirical auto-tuner has been used to address the neural network-based learning for performance comparison. In [43], a combination of stochastic differential equations and neural networks has been studied to extract the best combination of free parameters for the application of the economics dataset from Greater London using a Harris-Wilson model for a non-convex problem. In [44], a multi-label classification and complex regression problem has been addressed for auto-parameter tuning in deep learning models. In [45], ANNs have been used to predict the parameters for DE using 24 test problems from a Black-Box Optimisation Benchmarking dataset. In [46], parameter independent DE for analytic continuation has been studied using imaginary correlation functions of time. The parameters are embedded into the vectors which need to be optimised through the evolution. A study in [47] has proposed parameter optimisation for DE for the CED05 contest dataset that includes 25 complex mathematical functions with dimensions as high as 30.

## 3. Background

This section discusses the background techniques of differential evolution and artificial neural networks used in this research.

### 3.1 Differential Evolution (DE)

It is an efficient metaheuristic algorithm for numerical optimisation in which the output cannot be precisely defined from the input variables. For a given dimension of the data vector (string for example) it takes few input parameters such as number of points searching for the solution (population NP), mutation factor (F) and crossover rate (CR). The algorithm has 4 phases of execution: initialisation of population particles, mutation, crossover and then selection [48] as shown in Figure 2. This whole computation is repeated for a specified number of iterations (also known as generations) or a constrained time frame. Initialisation is usually done randomly from the normal distribution in the search domain followed by mutation which means finding the best parents to yield the child particle. Afterwards, crossover yields the child particle by varying proportions of parent particle attributes. The final step is selection, which means to update the current particle, how to use the new child along with other randomly selected particles with tuneable proportions. The idea is to maximise the randomness to avoid getting locked in local extrema. In our study, the simplest mutation formula (DE/rand/1) is used which is defined in (1) below:

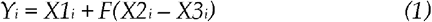

where *i* is the i^th^ point computed in an iteration from X1, X2 and X3 random points out of the population. Next, to increase the diversity of muted vectors, the crossover operation is performed which is defined in (2). It mixes the target vector with another random vector in the population in an adjustable proportion using random probability function which can be defined using CR value. CR rate thus defines the ratio in which the new trial vector *Ui* inherits the values from mutation vector.

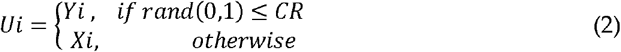

**Fig 2.**
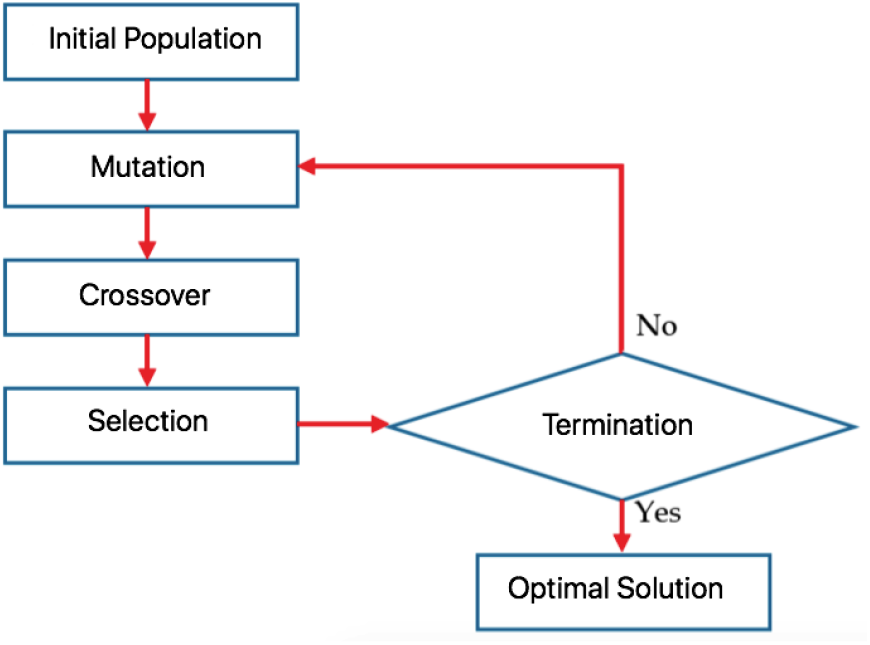
Flow of differential evolution metaheuristic algorithm

For the selection phase, the cost function of this new trial vector is calculated after the crossover phase to compare with the fitness of the target vector Xi. The better among the two is selected to update the army of points in the population. Let *f(*.) be the fitness function then, selection is defined as in (3) below:

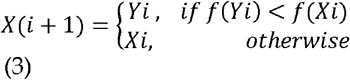

### 3.2 Artificial Neural Networks (ANN)

Artificial neural networks in data analysis got their origin from the behaviour of biological neurons on human brains. ANN consists of artificial nodes (neurons) which are interconnected through layers of other neurons to compute and refine the data using non-linear activation functions in the successive layers. Connection strengths among neurons is controlled by weight parameters (W). The simplest ANN in which we are interested is the multilayer perceptron (MLP), a three-layer structure is defined which contains the input layer, the hidden layer and the output. The standard structure of ANN is illustrated in Figure 3 and details can be found in [49].

**Fig 3.**
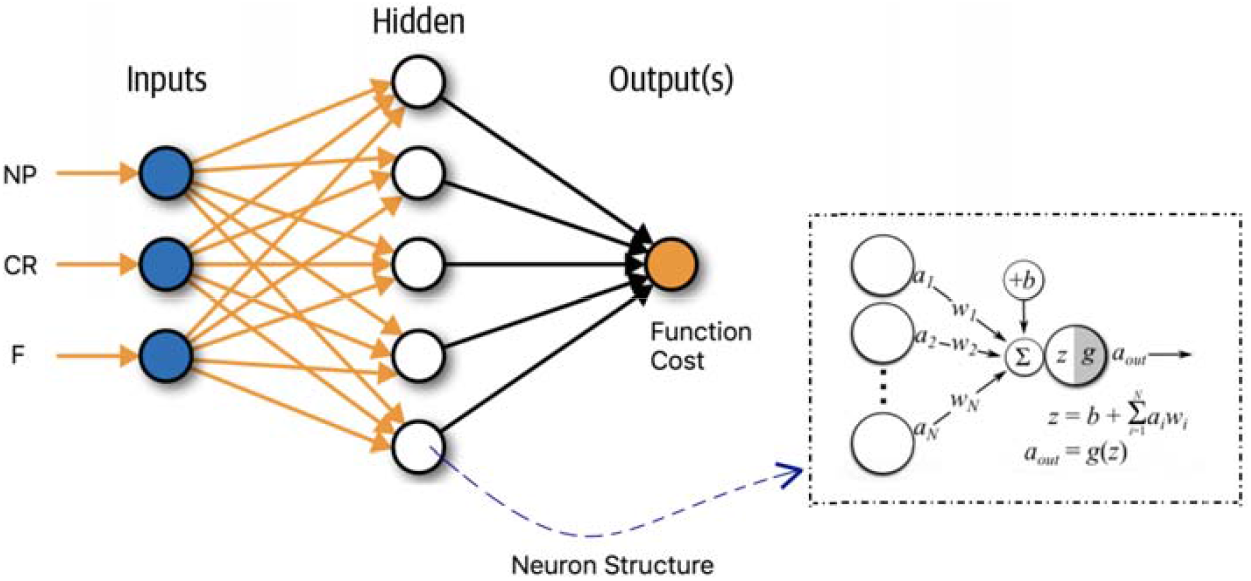
An example artificial neural network with three inputs, one hidden layer with five neurons and the output layer. The inside structure of each neuron has been shown in the dotted box. A node or neurons is the weighted average of the input signals added with a constant (bias) value and travels through a function which is usually non-linear in nature such as a tanh, sigmoid or ReLU (Rectified Linear Unit) function.

## 4. Results

Classical differential evolution uses three hyperparameters: the population size NP, a mutation factor F controlling the mutation rate, and a parameter CR that determines the extent of the uniform crossover [20] [26]operator used. We start with standard and fixed (non-adaptive) values of NP = 50, F = 0.4, CR = 0.1 as recommended by Storn and Price. NP is usually scaled to the length of the vector (number of input variables) to be optimised, but not much is known for problems that have a great many input variables [50], for instance directed protein evolution [51,52]; others are reviewed e.g. in [45].

### 4.1 Understanding the landscape ruggedness

To understand the nature of the landscape, it is convenient to create a random landscape in which, initially each element of the vector of length N=50 is set randomly to A to Z (Table 1). The random letters in each of the 10 vectors are the population candidates to search for the best rugged string of length 50.

**Table 1.**
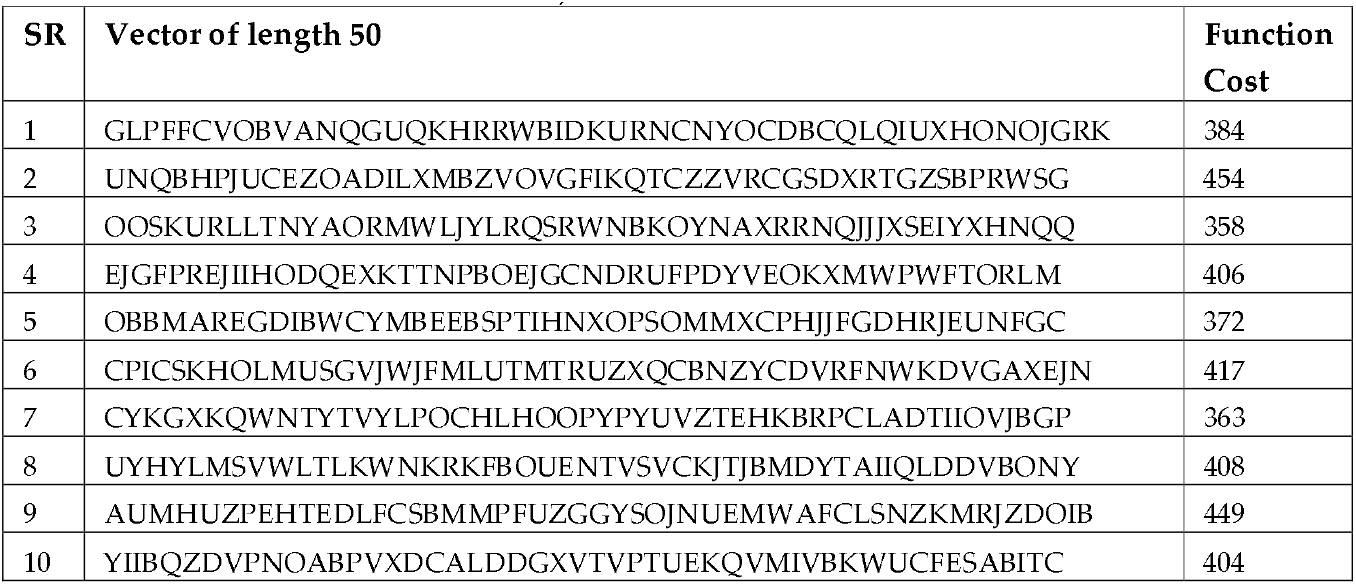
Random landscape of 10 vectors (population size) with dimensions = 50 (string length) and initial cost function value (summed consecutive differences)

After 10 iterations of the DE algorithm, the best particle and cost are as follows. Particle = APGZ TTTA PZIZ HZAZ ZDAZ BAEA ZAAZ AVWZ ABAZ AZAZ AZZZ AZAZ AZ, Cost function = 773, Benchmark = (26-1) × (50-1) = 1,225. Elapsed time is 0.1955 seconds. The values of Dimensions = 50, Np = 10, Population Crossover Rate (CR) = 0.8 and Mutation Factor (F) = 0.85 for this example. Figure 4 shows the consistent improvement in the cost function.

**Fig 4:**
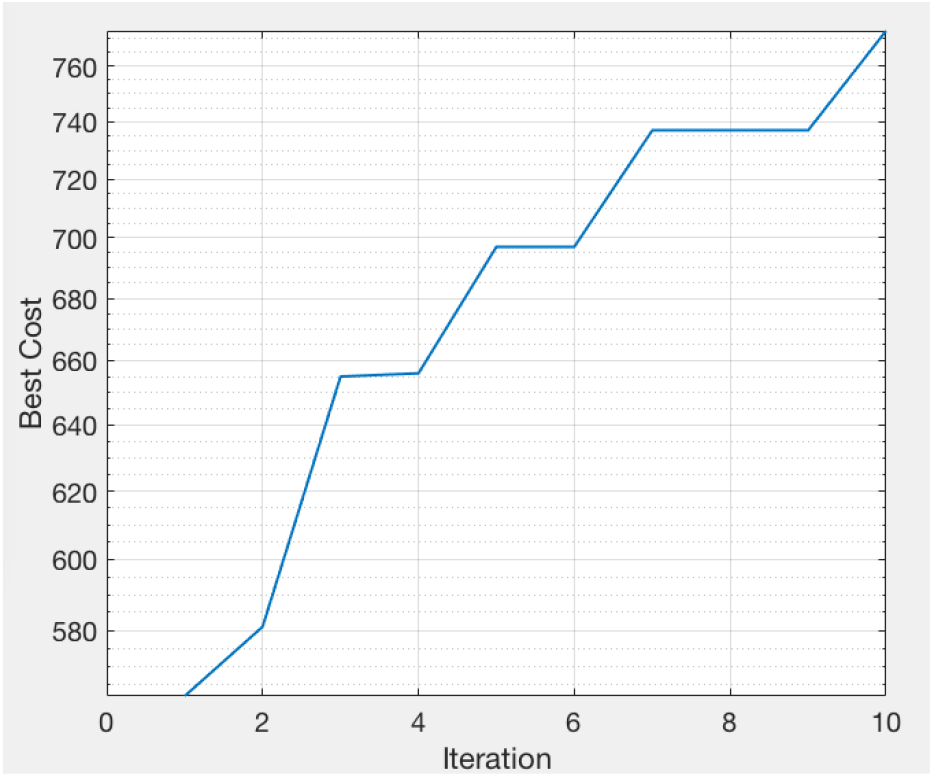
Cost function values for 10 iterations for string length = 50. The values of Dimensions = 50, Np = 10, Population Crossover Rate (PCr) = 0.8 and Mutation Factor (F) = 0.85.

### 4.2 Varying the hyperparameters in standard differential evolution

All simulations are performed in MATLAB 2018b software with a system configuration of MAC Air (2017), 1.8 GHz Intel Core i5 processor, 8GB 1600 MHz DDR3 memory, HD Graphics 6000 1536 MB and macOS version 10.13.6. The experiment was repeated for 500-dimensional string, 10-trials and 50 generations with NP in [50,500], CR in [0.1,1], F in [0.4,0.9]. The stats were recorded for the median and maximum function values for 10-k trials listed in Table 2. A total of 600 values were collected for various combinations of NP, F and CR values suggested in [45]. Out of those 600 values, the best 5 were selected and they were calculated for extended iterations/generations as shown in Table 3. It infers that the cost function value generally tends to increase with iterations but not always. This is because at higher dimensions the problem is highly difficult to solve and not always yields the best solution during the metaheuristic search.

**Table 2.**
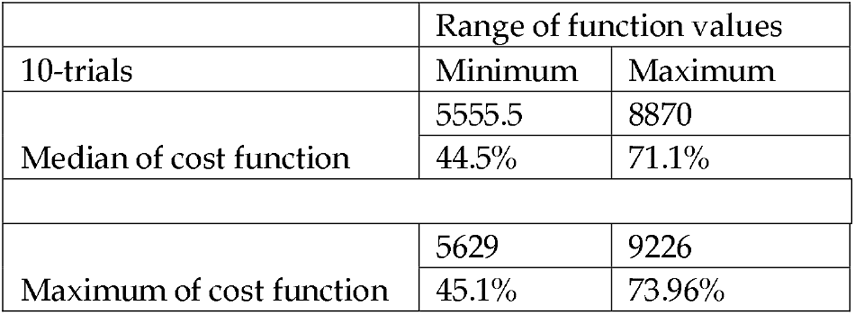
Benchmark value for 500-dimensional string is (26-1) × (500-1) = 12,475. NP = {50,100,…,500}, CR = {0.1,0.2,…1}, F = {0.4,0.5,…0.9} so total 10×10×6 = 600 experiment values. The min and max values refer to the range of cost function values obtained through the experiments.

**Table 3.** Top 5 cases among 600 trials analysed for increased number of iterations from 100 to 1000. The winners are highlighted in bold text.

| CASE1 | NP | F | CR | Iterations | 100 | 200 | 300 | 500 | 1000 |
| --- | --- | --- | --- | --- | --- | --- | --- | --- | --- |
|  | 500 | 0.9 | 1 | Median | 9632 | 10300 | 10375 | 10300 | **10375** |
|  |  |  |  | Maximum | 9716 | 10550 | 10425 | 10375 | **10625** |
| CASE2 | NP | F | CR | Median | 7935 | 8970 | 9269 | 9280 | **9295** |
|  | 500 | 0.8 | 0.9 | Maximum | 8222 | 9106 | 9322 | **9592** | 9538 |
| CASE3 | NP | F | CR | Median | 8801 | 9246 | 9159 | 9012 | **9230** |
|  | 500 | 0.9 | 1 | Maximum | 8901 | 9362 | 9342 | 9181 | **9328** |
| CASE4 | NP | F | CR | Median | 7002 | 9037 | 9218 | **9536** | 9242 |
|  | 500 | 0.8 | 0.9 | Maximum | 7862 | 9152 | 9529 | **9677** | 9651 |
| CASE5 | NP | F | CR | Median | 6075 | 8211 | 8500 | **8675** | 8450 |
|  | 500 | 0.8 | 0.9 | Maximum | 6250 | 8451 | 8500 | **8775** | 8500 |

**Table 4** and **Figure 5** illustrate the normalised function values for various combinations of population size and generations. For median of cost function values, it has been found that 9196 value is obtained with the NP=50 and Gen=500. For the maximum cost function values the value 9373 is found when NP=125 and Gen=200. Thus, it is not easy to say which of the NP and Generation values are the best ones to yield the best function cost but generally more generations with bigger population size is a good combination.

**Table 4.** Normalised comparison (NP*Gen constant) of cost function values (median and maximum over 10-trials) for F=CR=0.9, string length = 500. The values of VP and generations have been varied such that the product of NP and Gen is 25000. Figure 5 is the visual illustration.

| Sr. | NP | Generations | Median | Maximum |
| --- | --- | --- | --- | --- |
| 1 | 500 | 50 | 8236 | 8272 |
| 2 | 250 | 100 | 9109 | 9394 |
| 3 | 125 | 200 | 9150 | **9373** |
| 4 | 50 | 500 | **9196** | 9304 |
| 5 | 10 | 2500 | 8137 | 8601 |

**Fig 5.**
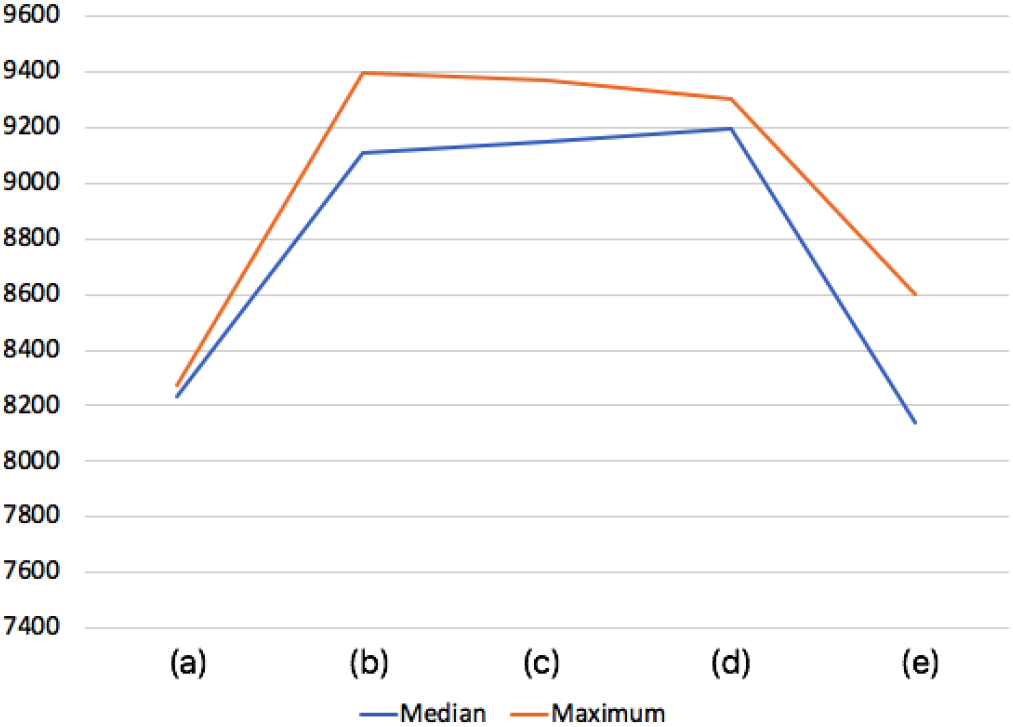
Normalised comparison of various values of function cost for median and maximum using F=0.9, CR=0.9, string length (dimension)=500, 10-trials. (a) NP=500, Gen=50 (b) NP=250, Gen = 100 (c) NP=125, Gen=200 (d) NP=50, Gen=500 (e) NP=10, Gen=2500. The values of NP and Generations have been chosen such that the product remains constant (=25000) to analyse the effect of reducing population and increasing generations.

**Tables 5-7** illustrate time consumption analysis against variations in NP, string length and iterations of calculations. **Figures 6-8** are the graphical representations of these tables for the ease of visualisation. The time variations are almost linear giving the idea about the problem complexity in regard to the variable changes.

**Table 5.** For 50 iterations, string length 500 (data vector dimension), F=0.4, CR = 0.1 the time in seconds for various NP values has been calculated ranging from 50 to 500 for 10-trial experiments. **Figure 7** is the graphical rendering of this table.

|  |  |  |  |  |  |  |  |  |  |  |
| --- | --- | --- | --- | --- | --- | --- | --- | --- | --- | --- |
| NP | 50 | 100 | 150 | 200 | 250 | 300 | 350 | 400 | 450 | 500 |
| Time | 1.32 | 2.46 | 3.73 | 4.88 | 6.24 | 7.43 | 8.94 | 10.32 | 11.71 | 14.07 |

**Table 6.** String length (Dimension) versus time consumption in seconds for fixed values of NP=100, F=0.4, CR=0.1, iterations = 50 and 10-trials for experiments to calculate median and mean function costs. The time consumption varies almost linearly with the string length. **Figure 8** shows the graphical representation

|  |  |  |  |  |  |  |  |  |  |  |
| --- | --- | --- | --- | --- | --- | --- | --- | --- | --- | --- |
| Dim | 100 | 200 | 300 | 400 | 500 | 600 | 700 | 800 | 900 | 1000 |
| Time | 0.7992 | 1.1327 | 1.7634 | 2.1091 | 2.5229 | 2.8815 | 3.2637 | 3.5849 | 3.9694 | 4.2432 |

**Table 7.** Iterations and time in seconds comparison for fixed values of string length = 500, NP=100, F=0.4, CR=0.1 and 10-trial experiments. The relation is almost linear and is rendered in **Figure 9**.

|  |  |  |  |  |  |  |  |  |  |  |
| --- | --- | --- | --- | --- | --- | --- | --- | --- | --- | --- |
| Gen | 50 | 100 | 150 | 200 | 250 | 300 | 350 | 400 | 450 | 500 |
| Time | 2.56 | 4.75 | 7.19 | 9.99 | 11.64 | 13.99 | 16.49 | 20.28 | 20.99 | 25.55 |

**Fig 6.**
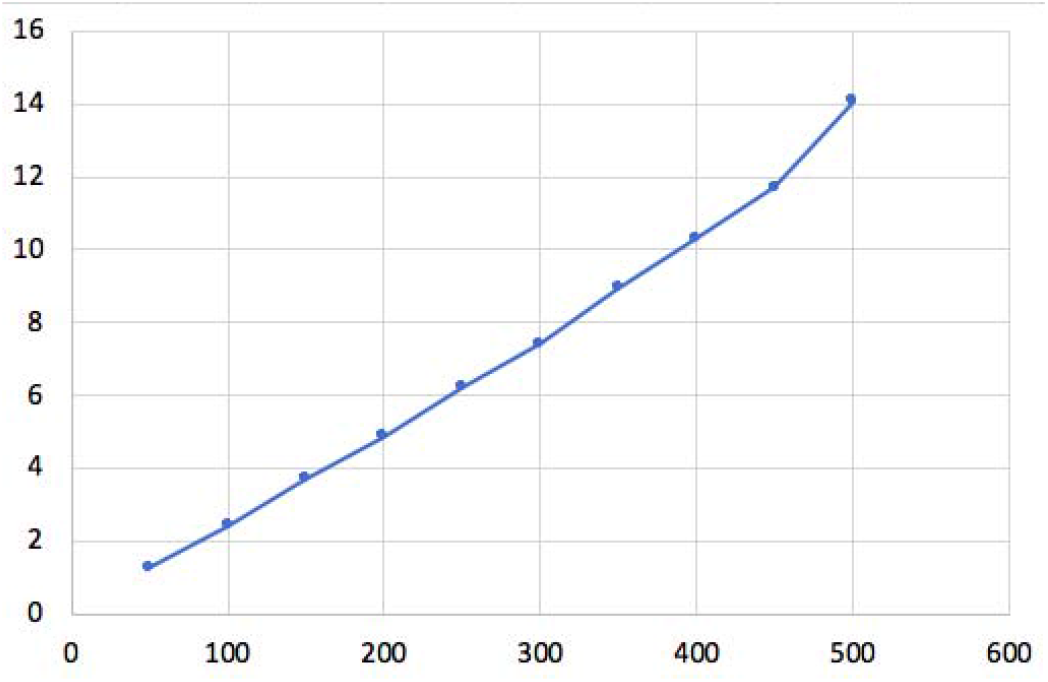
Graph between NP (x-axis) and Time in seconds (y-axis) for fixed values of string length = 500, F = 0.4, CR = 0.1 and iterations = 50 for 10-trial experiments. The time consumption is nearly linear with the population size.

**Fig 7.**
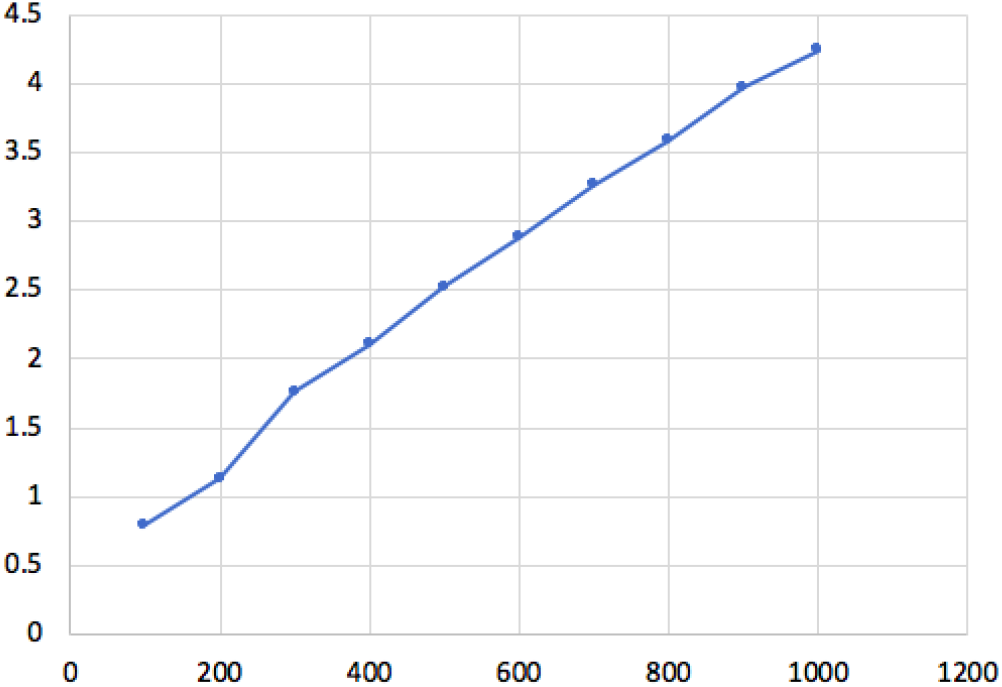
Graph between string length (x-axis) and Time in seconds (y-axis) for fixed values o NP=100, F=0.4, CR=0.1, iterations = 50 and 10-trials for experiments to calculate median and mean. The time consumption varies almost linearly with the string length.

**Fig 8.**
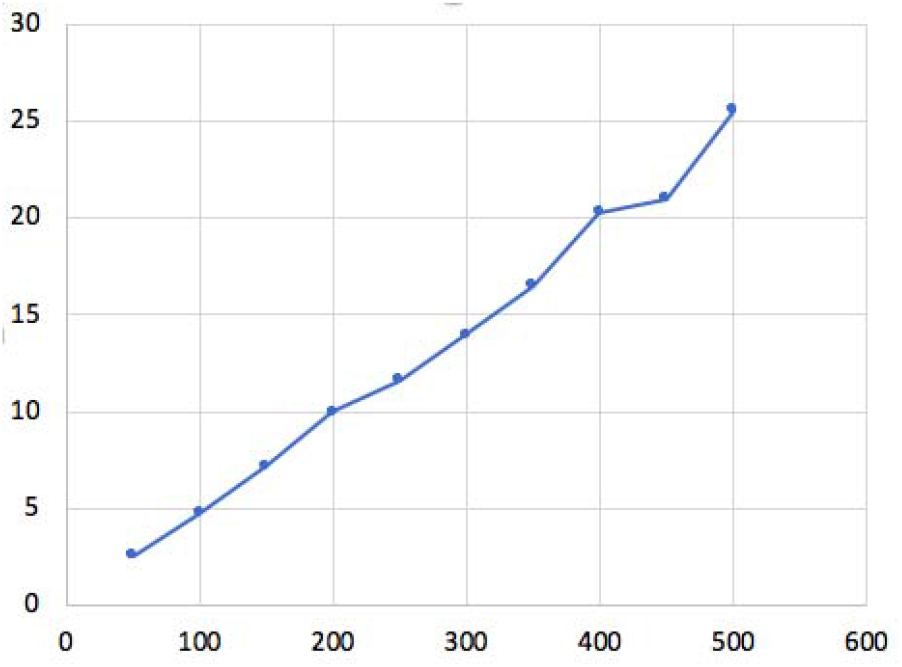
Graph between iterations (x-axis) and time in seconds (y-axis) for fixed values of string length = 500, NP=100, F=0.4, CR=0.1 and 10-trial experiments. The relation is almost linear.

### 4.3 Training Data and ANN Training for Automatic Parameter Tuning

The neural network has been trained for 600 data points which includes three independent input variables: Population (NP), Mutation Factor (F) and Cross-mutation Rate (CR). The values range for NP is [50,500], F is [0.4,0.9], CR is [0.1,1] chosen according to our experiments and suggestion given in [40]. These 600 training data points have been collected by running 10-fold trials for each entry to find out median and maximum cost function values and is quite a time consuming task. This is due to finding the right range of these 3 hyperparameter values and then running the code to compile the data for few hours. So the differential evolution function is called 600×10 = 6,000 times for various values of these three hyperparameters. The training sample data and the computed median and maximum function costs have been rendered in **Table 8**. Afterwards, Neural Network Fitting in MATLAB has been used for regression using three input variables and two target variables respectively. The default values of the model have been used, for example ratio of training: validation: testing is 70:15:15, number of hidden neurons = 10, Bayesian Regularization training algorithm for training. **Figures 9-10** demonstrate the frequency histogram of error values in relation to instances. After the training of ANN, the error of actual versus predicted values were collected for all training and tested values among 600 data points and plotted to see the quality of the trained model. Most of the error values are 0 which indicates that the model is well generalised and validated.

**Table 8.**
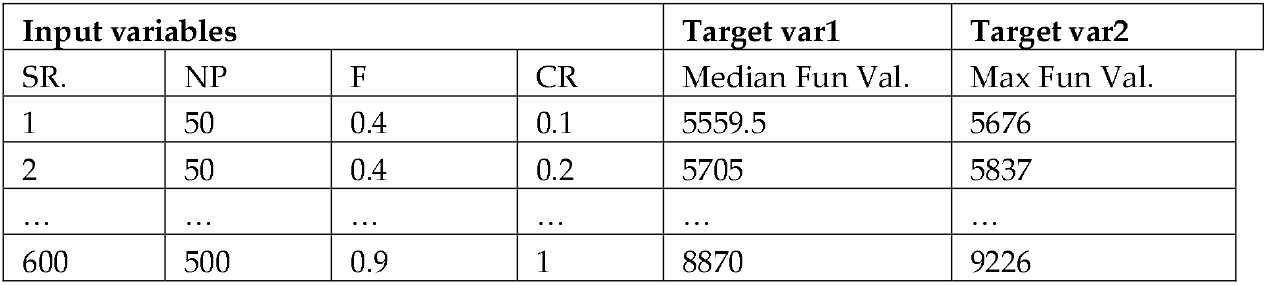
Training data for ANN containing 3 input variables and two target variables. The NP ∈ {50,100,…,500}, F ∈ {0.4,0.5,…,0.9}, CR ∈ in {0.1,0.2,…,1} so a total of 10×6×10 = 600.

**Fig 9.**
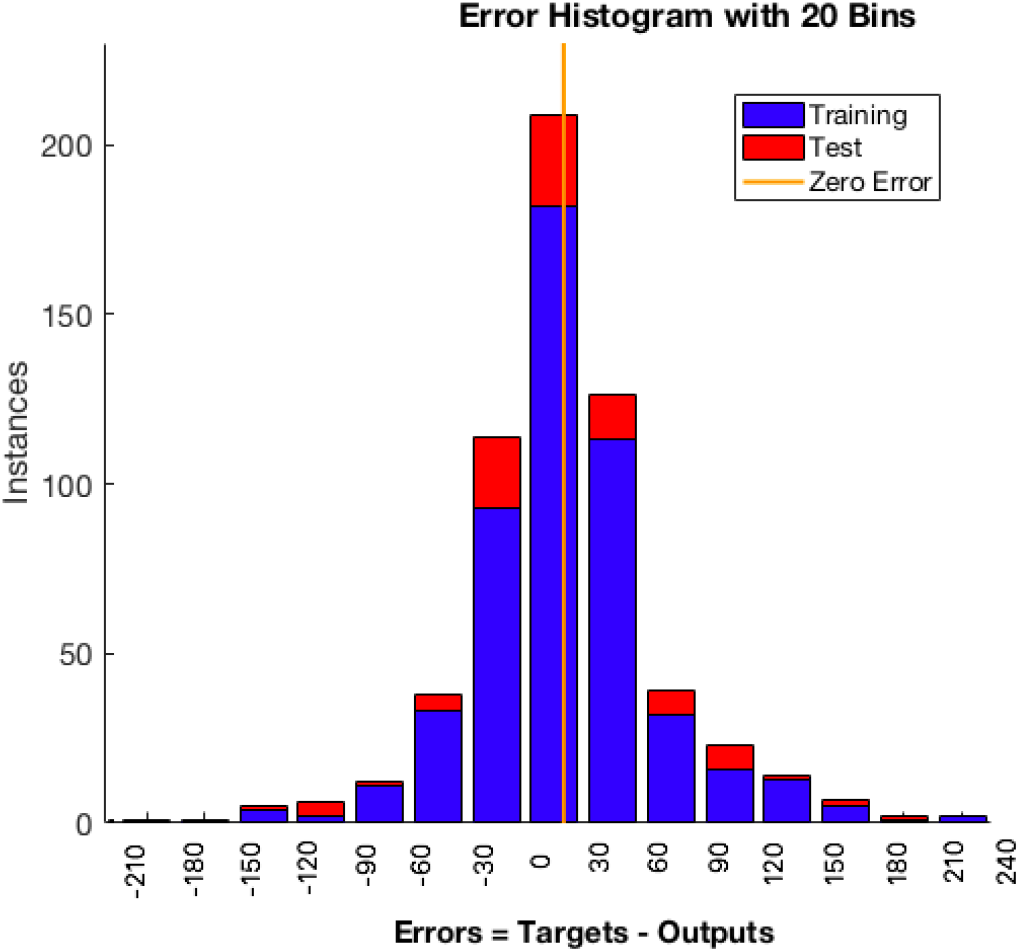
Error histograms for training, testing and validation for 600 examples after ANN training for *median* cost function values. Ratio of train:validate:test used is 70:15:15. The target function used is the median cost function value from 10-trial experiments and input values to ANN were NP, F and CR.

**Fig 10.**
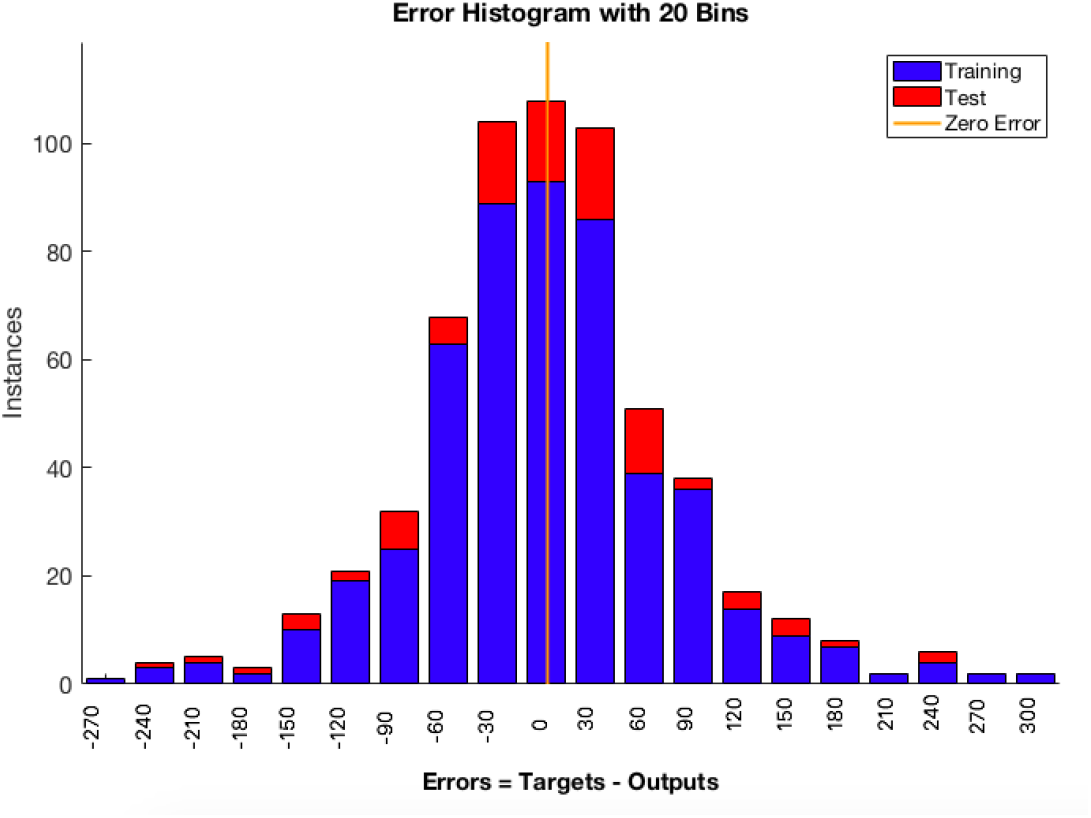
Error histograms for training, testing and validation for 600 examples after ANN training for *max* cost function values. Ratio of train:validate:test used is 70:15:15. The target function used is the maximm cost function value from 10-trial experiments and input values to ANN were NP, F and CR.

**Tables 9-10** show the function costs on 10-trial average for the mean squared error (MSE) and regression coefficient (R) values as the number of hidden neurons increase during the ANN training.

**Table 9.** For the median cost function values (from 10-trials) the following are the MSE and R values for increasing number of neurons in ANN training

| Median<br>Function<br>Cost |  | Neurons | 10 | 20 | 30 | 40 | 50 | 60 | 70 | 80 |
| --- | --- | --- | --- | --- | --- | --- | --- | --- | --- | --- |
|  | Training | MSE | 3008.8 | 2102.2 | 1264.8 | 1472.8 | 1572.3 | **1108.8** | 1653 | 1540.5 |
|  |  | R | 0.9975 | 0.9983 | 0.9987 | 0.9990 | 0.9985 | 0.9988 | **0.9989** | **0.9989** |
|  | Testing | MSE | **2858** | 3624 | 5883.7 | 5616.6 | 4622.5 | 7715.5 | 3162 | 5431.8 |
|  |  | R | 0.9948 | 0.9959 | 0.9947 | 0.9962 | 0.9974 | 0.9946 | 0.9936 | 0.9957 |

**Table 10.** For the maximum cost function values (from 10-trials) the following are the MSE and R values for increasing number of neurons in ANN training

| Max<br>Function<br>Cost |  | Neurons | 10 | 20 | 30 | 40 | 50 | 60 | 70 | 80 |
| --- | --- | --- | --- | --- | --- | --- | --- | --- | --- | --- |
|  | Training | MSE | 8738 | 6682.7 | 6021.4 | 5308.3 | 5790.4 | 5791.7 | 5557.3 | **5059.4** |
|  |  | R | 0.9937 | 0.9954 | 0.9958 | 0.9963 | 0.9958 | 0.9960 | 0.9961 | **0.9964** |
|  | Testing | MSE | 1148.3 | **1192.8** | 1308.1 | 1228.3 | 2106.7 | 1262.8 | 1689.5 | 1252.9 |
|  |  | R | **0.9934** | 0.9916 | 0.9912 | 0.9914 | 0.9872 | 0.9908 | 0.9877 | 0.9926 |

**Figures 11-12** show typical curves of the median and maximum fitness as a function of the number of evaluations for 10-trials. A 4-d representations of the test results obtained by ANN with 3,971 points has been shown, for a range of NP ∈ {50,75,…,500}, F ∈ {0.4,0.45,…,0.9}, CR ∈ {0.1,0.15,…,1} values. The colour illustrates the range of cost function values predicted by ANN. Yellow means higher (better) function value and blue means lower cost function value for that combination of NP, F and CR hyper-parameters. It can be observed that the best function values (both maximum and median values) are obtained from higher NP, CR and F values in general. It can be seen that the best function costs correspond to maximum values of NP, F and CR values for both figures.

**Fig 11.**
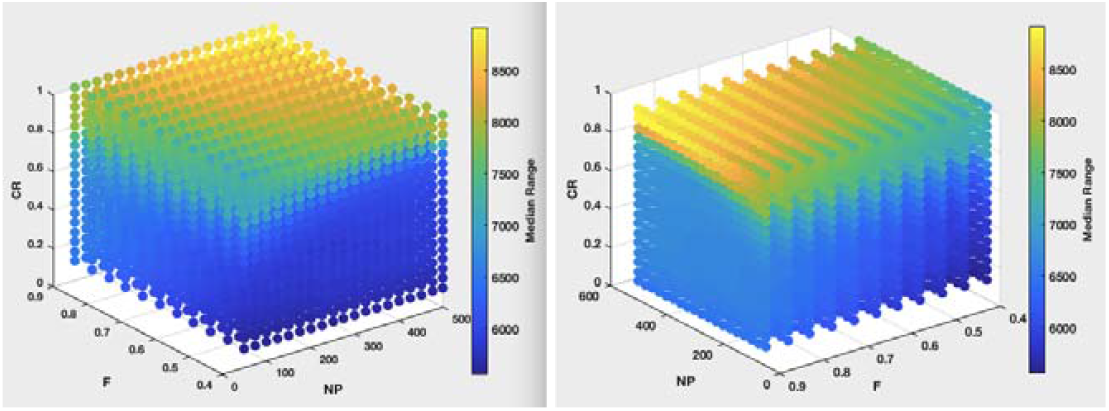
The demonstration of 3,971 test points (19×11×19) parameter combinations of NP ∈ {50,75,…,500}, F ∈ {0.4,0.45,…,0.9}, CR ∈ {0.1,0.15,…,1}. The colour depicts the median of cost function values of 10-trial runs. The yellow shows higher (better) values and blue shows lower cost function values for those combinations of NP, F and CR. Left and right graphs are two perspective view of the same 3-d figure.

**Fig 12.**
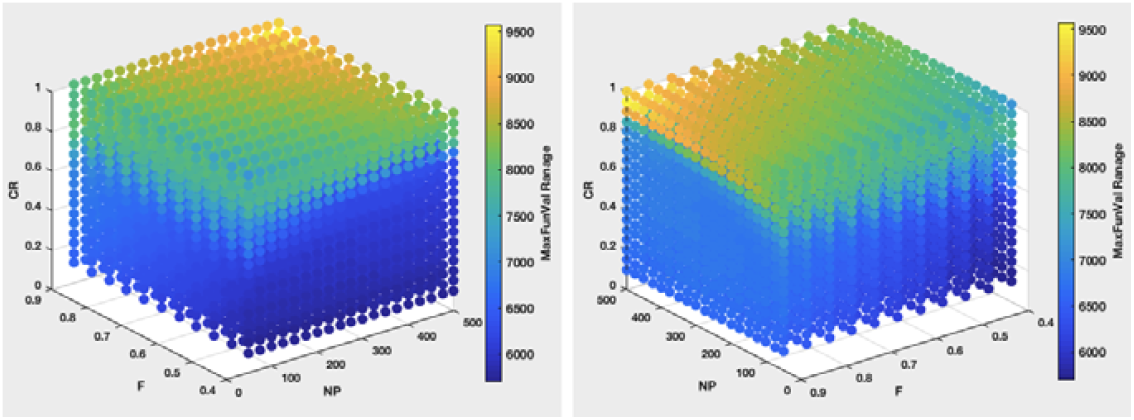
The demonstration of 3,971 test points (19×11×19) parameter combinations of NP ∈ {50,75,…,500}, F ∈ {0.4,0.45,…,0.9}, CR ∈ {0.1,0.15,…,1}. The colour depicts the maximum value of cost function for 10-trial runs. The yellow shows higher (better) values and blue shows lower cost function values for those combinations of NP, F and CR. Left and right graphs are two perspective view of the same 3-d figure.

### 4.4 Validation

Among the total tested values (3,971) we chose the best 10% values with maximum cost function values (yellow in Figure 11-12) to calculate the actual cost functions through differential evolution on the average of 10-trials. Afterwards, both the continuous variables (ANN suggested output and actual DE values) for (NP, F, CR) hyper-parameter combinations have been analysed using Pearson’s correlation coefficients (ρ). For both cases of DE versus ANN test results ρ_MEDIAN_ = 0.7 and ρ_MAXIMUM_ = 0.68. It indicates a significant positive relationship among two variables [53]. Thus the hyperparameter optimisation in differential evolution with summed local difference strings can be performed efficiently using the neural network simulation.

### 4.5 Comparison with other state-of-art

To analyse the effectiveness of ANN-based parameter tuning of DE, we compared the following techniques: (i) Grid Search, (ii) Random Search, (iii) Sequential Model-based Algorithm Configuration (SMAC) [44], (iv) Optuna [41] (v) ANN in **Table 11** and **Figure 13**.

**Table 11.** Hyperparameter optimisation for differential evolution algorithm to search for {NP,F,CR} by various algorithms. Benchmark value for string of length (=NP) of 500 = (26-1)*(500-1)=12,475 and 10 trial runs. Generations for DE=100 for all of them consistently and max of 10 trials experiments for the best cost calculations. Accuracy is the ratio of best cost and the benchmark (=12,475).

| Methods | Details | Hyperpram Values | Best Cost | Accuracy |
| --- | --- | --- | --- | --- |
| Grid Search | NP$\in$[50,500],<br>F=CR=[0.40, 0.41,...1.0} | NP=390, F=0.95,<br>CR=0.90 | 8616 | 0.6907 |
| Random Search | NP$\in$[50,500],<br>F, CR $\in$ [0.4,1] | NP=450, F=0.91,<br>CR=0.88 | 8572 | 0.6871 |
| SMAC | NP$\in$[50,500],<br>F, CR $\in$ [0.4,1] | NP=410, F=1, CR=0.98 | 8990 | 0.7206 |
| Optuna | NP$\in$[50,500],<br>F, CR $\in$ [0.4,1] | NP=430, F=,0.87<br>CR=0.92 | 9244 | 0.7410 |
| ANN | NP$\in$[50,500],<br>F=CR=[0.40, 0.41,...1.0} | NP=500, F=0.90,<br>CR=0.89 | 9438 | 0.7566 |

**Fig 13.**
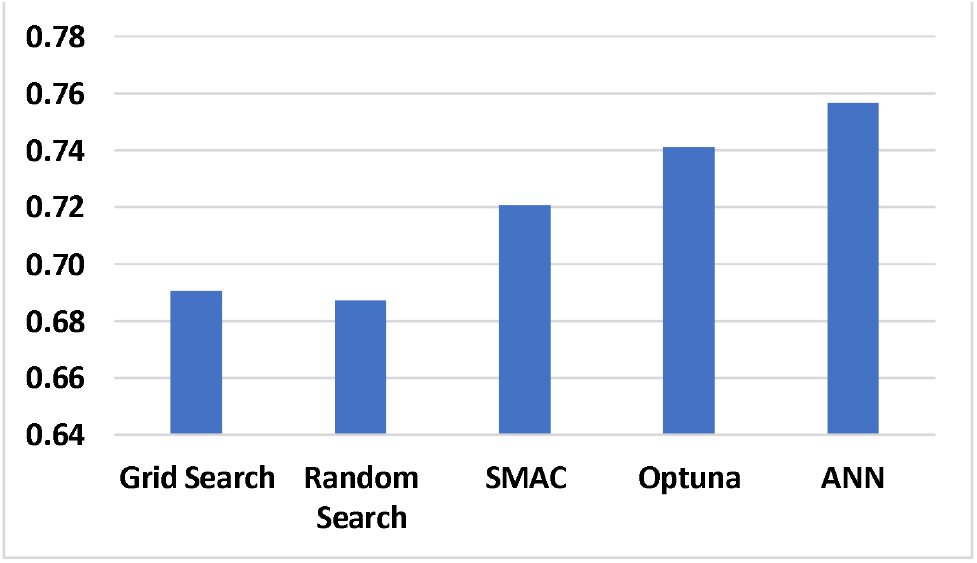
Graphical representation of accuracy comparision of Table 11. Our ANN performed better than the other state-of-art hyperparameter techniques studied.

Grid Search suffers from the curse of dimensionality, resulting in an explosion in the number of possible evaluations, which is improved by Random Search. But random search is not well sorted and may miss the potential extremas. SMAC uses random forests (RF) and is a Bayesian optimiser in which RF helps in categorical variables to support large search space hyperparameter searches and is well scalable for increasing the number of training samples. It is available online https://github.com/automl/SMAC3. Optuna [41] is recent software used for hyperparameter optimisation using define-by-run API, pruning and search strategy implementation, versatile utility including distributed computing, scaling and interactive interface for users to modify the search space parameters dynamically. It is available online https://github.com/optuna/. Hyperparameter auto-tuning has also been performed for sparse Bayesian learning (SBL) in [42] using neural network-based learning, and has shown considerable improvement in recovery performance and convergence rate.

ANN performed better compared to Grid Search, Random Search, SMAC and Optuna (**Table 11 and Fig 13**.), but requires a considerable amount of time to generate the training data to simulate the behavior of the rigged function outcome for given values of NP, F and CR. Afterwards it is able to predict the new values of hyperparameters to search for the optimal combination of tuning parameters. ANN requires much less time as compared to DE to calculate the cost function once it is trained on a good sized set. For the sake of simplicity, only 600 examples have been used to train ANN for this case study, but a few thousand runs of DE would be better to yield the training data for ANN.

## 5. Conclusion

In this research, hyperparameter optimisation in differential evolution has been studied using Summed Local Difference Strings, which is a rugged but easily calculated landscape for combinatorial search problems with wide applicability in numerical optimisation, biological sciences, finance and organisational management. Differential evolution is a powerful numerical optimisation technique for non-differentiable and complex functions which cannot be nicely defined in mathematics, but it has three hyperparameters (NP, F, CR) to be optimised. In this study a machine learning technique has been exploited to suggest the best possible combinations of hyper-parameters instead of a tedious grid search. The limitation of the technique is that training data collection is time consuming and needs a careful analysis of input variable ranges (NP, F, CR). Two output variables were recorded (median and maximum value) after 10-trial experiments of each combination of the hyperparameters. Finally, testing of the machine learning model has been employed on a bigger data (3,971) and the top 10% test results were compared with the actual DE results yielding a Pearson correlation coefficient of ∼0.7. Specifically, it was found that larger values of NP, F and CR hyper-parameters yield better outcomes.

In future, we plan to explore more bio-inspired algorithms and other ways to search for hyperparameters such as to embed the hyper-parameters in the very population being optimised. We will also explore binary strings and real valued strings with an extended range of values beyond 26 with advanced options of mutation and cross-over equations.

## Acknowledgements

We thank the UK EPSRC and AkzoNobel for financial support via the SusCoRD project, grant EP/S004963/1. DBK thanks the Novo Nordisk Foundation for financial support (grant NNF20CC0035580).

